# Vaccine-induced and hybrid immunity against SARS-CoV-2 variants of concern in two cohorts in Queensland, Australia (2021-2022)

**DOI:** 10.1101/2024.11.11.623100

**Authors:** Rhys H. Parry, Yingjun Shang, Julian D.J. Sng, Christopher L. D. McMillan, Naphak Modhiran, Alberto A. Amarilla, Ariel Isaacs, Benjamin Liang, David A. Muller, Daniel Rawle, Andreas Suhrbier, Stevie Anderson, Samuel Kelly, Daniel Watterson, Lucy D. Burr, Alexander A. Khromykh

**Author notes:** These authors contributed equally (Rhys H. Parry, Yingjun Shang).

## Abstract

The spike glycoprotein of severe acute respiratory syndrome coronavirus 2 (SARS-CoV-2) is the main target for vaccine development, as antibodies generated against the spike protein are the most immunodominant and neutralizing against the virus. However, variants of concern (VOC), often containing multiple mutations within neutralizing epitopes, confer immune evasion of the response generated by current SARS-CoV-2 vaccines. To assess the immunogenicity and virus-neutralisation ability of antibodies induced by individual or combination of Vaxzevria (AstraZeneca: AZ) and Comirnaty (Pfizer: PZ) vaccines and of a hybrid immunity induced in people who were also infected with SARS-CoV-2, longitudinal sera samples of participants recruited through the David Serisier Research biobank (Mater Research Hospital) or at the University of Queensland were collected. ELISA with a panel of purified Spike proteins from ancestral, alpha, delta and omicron BA.1 and BA.2 VOCs showed significantly (by ∼2 to 6-fold) reduced IgG antibody titres against Spike proteins from Omicron BA.1 and BA.2 compared to ancestral strain regardless of the number of vaccinations or presence of infection. Neutralisation assays showed reduced activity against delta and omicron BA.1, BA.5 and BA.5 VOCs. However, the differences were in general less pronounced than in the ELISA assay and some were not statistically significant, particularly after four (two AZ and two PZ) vaccinations. We also generated by circular polymerase extension reaction an attenuated SARS-CoV-2 strain with deletion of all accessory genes, ORF 3, 6, 7 and 8, based on the ancestral (QLD02) virus backbone (QLD02_Δ3678_) and validated it in virus-neutralization assays with our panel of sera samples. We showed the attenuation of the QLD02_Δ3678_ virus in Vero E6 and human Caco2 cells. We demonstrated that neutralization assays with the wild-type QLD02 virus and QLD02_Δ3678_ virus were concordant, providing a safe platform for neutralisation assays in BSL2/PC2 settings.

## Introduction

The COVID-19 pandemic, caused by the SARS-CoV-2 (Name: *Severe acute respiratory syndrome-related coronavirus*, Species Betacoronavirus pandemicum, Genus: *Betacoronavirus*), has highlighted the importance of monitoring adaptive immunity elicited from vaccination and natural infection to assess the duration of protection against SARS-CoV-2. Longitudinal monitoring of individuals allows to assess the duration of protection against SARS-CoV-2, compare the durability of different vaccines, track the dynamics of waning antibody titers, and identify the emergence of neutralization-resistant variants of concern.

Following SARS-CoV-2 infection or vaccination, the immune system mounts a robust antibody response, generating IgG, IgA, and IgM immunoglobulins. Seroconversion typically occurs within 15 days post-infection, with IgM and IgG peaking around 18 and 20 days, respectively^1,2^. Spike-specific antibodies typically neutralize the virus by binding to the receptor-binding domain (RBD), and block its interaction with cell surface receptor proteins, as well as mediating antibody-dependent cellular cytotoxicity and phagocytosis through Fc receptors on immune cells.

In Australia, Pfizer/BioNTech’s BNT162b2 mRNA vaccine (PZ) formulated as RNA–lipid nanoparticles of nucleoside-modified mRNA^3^ and the Oxford/AstraZeneca’s ChAdOx1-S non-replicating adenovirus-vectored vaccine (AZ)^4^ were the first approved COVID-19 vaccines by the Therapeutics Goods Administration (TGA). Both vaccines express the SARS-CoV-2 spike protein, and primary administration involves two doses of AZ vaccine (AZ_1_/AZ_2_) with an 8-12-week interval, and in some individuals followed by two doses of PZ vaccine (PZ_1_/PZ_2_) administered 4-8 weeks apart. Peak antibody titres are observed within 2-4 weeks, followed by a log-linear decline over 4 to 16 weeks^5–7^. The second dose significantly boosts antibody responses, leading to higher overall antibody titers^8^. Factors such as, age, sex, infection severity and hospitalisation status influence the peak magnitude of the IgG response^5,9–11^.

While the primary regimen of AZ and PZ vaccines confer strong immunogenicity and higher short-term efficacy, their effectiveness wanes considerably 5-8 months post-vaccination. Despite this, they maintain effective real-world protection against hospitalisation and death after 6 months with 79% and 90% efficacy, respectively^12^. Although absolute IgG titres against the ancestral spike protein decline over time, durable, long-lived neutralising antibodies, particularly those targeting the RBD, exhibit a sustained plateau phase^2,13^.

Variants of concern (VOC) are strains of SARS-CoV-2 that have undergone multiple mutations, leading to changes in transmissibility, disease severity, or immune escape potential^14^. The first VOC, Alpha (B.1.1.7), identified in England, was estimated to have a 40–80% increased transmissibility than ancestral SARS-CoV-2^15^. Key defining mutations in the spike protein (D614G and ΔH69/V70) contribute to this phenotype^16,17^. The Delta VOC (B.1.617.2) demonstrated significant immune evasion with several studies reporting between 3.2-to 5-fold reduction in neutralisation with a primary AZ or PZ regimen^18–20^. The Omicron variant (B.1.1.529) includes several sublineages (BA.1, BA.2, BA.5) that are the most immunoevasive, with more than 15 mutations on average in the RBD. The neutralizing activity of sera from convalescent and double-vaccinated individuals was undetectable or very low against Omicron variants compared with the ancestral virus^21,22^, whereas neutralizing activity of sera from individuals who had been exposed to spike three or four times through infection and/or vaccination was maintained, although at significantly reduced levels^22^. It was later shown that Omicron BA.1 escapes the majority of existing SARS-CoV-2 neutralizing antibodies^21^.

The overall reduction in circulating IgG against the Spike protein and the emergence of immunoevasive VOCs suggested that the initial primary two-dose regimen was becoming less effective overall against VOC with less than 20% effectiveness against laboratory-confirmed Omicron infection at 6 months^23^. To counteract waning immunity and reduced neutralisation of VOCs, booster doses have been introduced^24^. These booster doses, typically administered six months after the primary vaccination series, have been shown to significantly enhance antibody levels and improve protection against emerging VOCs, although 9 months after booster administration vaccine effectiveness was less than 30% against laboratory-confirmed infection and symptomatic disease^23^. In Australia, the TGA approved the Pfizer COVID-19 vaccine on October 27, 2021 after interim results showed up to 95.3% vaccine efficacy^25^.

To evaluate the immunogenicity and virus-neutralizing efficacy of responses induced by vaccinations we sampled two cohorts in Brisbane, Australia, one containing health workers recruited through David Serisier Research biobank (Mater Research Hospital) and another one containing participants from the University of Queensland. Samples were collected throughout the study from the same participants following 2, 3 and 4 vaccinations with combinations of Astra Zeneca (AZ) and Pfizer (PZ) vaccines. A separate set of samples was collected from participants who were vaccinated and infected with SARS-CoV-2 to assess hybrid immunity. The IgG antibody titres were assessed by ELISA against purified Spike proteins from different VOCs and virus-neutralising antibody titers were assessed against different VOCs in high throughput microneutralizations assay. Towards the development of a safer virus neutralization assay that can be performed in BSL2 facilities, we generated an attenuated SARS-CoV-2 with deletion of all accessory genes and compared its efficacy in neutralization assay with that of full-length infectious virus.

## Materials and Methods

### Human ethics, sera and nasal swab collection and processing

The collection of human sera was carried out following the approval of the Mater Misericordiae Ltd Human Research Ethics Committee (Reference: HREC/MML/68320) or the University of Queensland Human Research Ethics Committee (References: 2021/HE000362, 2021/HE000139). Blood samples, approximately 5mL in volume, were collected in serum-separating tubes and processed by centrifugation at 1000g for 10 minutes at 4°C. After centrifugation, the sera were aliquoted into individual, single-use tubes and stored at -80°C. Nasopharyngeal aspirates of infected individuals were collected from consenting COVID-19-positive patients in Brisbane, Australia and collected for the recovery of SARS-CoV-2 isolates as previously described^26^.

### Cell lines

Vero E6 cells, derived from African green monkey kidney epithelia and Vero E6 stably expressing human TMPRSS2 (VE6-hTMPRSS2 Clone #1), as previously described^27^, were cultured at 37 °C and 5% CO_2_ in Dulbecco modified Eagle medium (DMEM; Gibco, Thermo Fisher, USA) supplemented with 10% fetal calf serum (FCS; Bovogen; USA), 1% GlutaMAX™ (Cat: 35050079, Thermo Fisher, USA), and 100 U/mL penicillin-streptomycin (Gibco, Thermo Fisher, USA), VE6-hTMPRSS2 cells were grown in the same media as Vero E6 cells with addition of 30µg/mL of puromycin dihydrochloride (A1113803, Thermo Fisher, USA). Caco2 (human colorectal adenocarcinoma cells) were cultured in RPMI Medium 1640 (Gibco, Thermo Fisher, USA) supplemented with 10% FCS and 100 U/mL penicillin-streptomycin. Human embryo kidney tissue cells stably expressing ACE2 and TMPRSS2 (HEK293T-ACE2-TMPRSS2) cells were generated by transduction of HEK293T-ACE2 cells, provided by Jesse Bloom (Fred Hutchinson Cancer Research Centre, Washington, USA) with a lentivirus containing puromycin-selectable codon-optimized human TMPRSS2 gene^28^ and validated with anti-TMPRSS2 antibody (ab109131, Abcam, UK). These cells were selected and propagated in the media supplemented with 2µg/mL puromycin dihydrochloride.

### SARS-CoV-2 viruses

This study used five low-passage SARS-CoV-2 viruses recovered from nasopharyngeal aspirates of infected individuals provided by the Queensland Health Forensic and Scientific Services, Queensland Department of Health, and the Queensland Institute of Medical Research. These strains included: I) an early ancestral (PANGO v.4.2 Lineage A) Australian isolate hCoV-19/Australia/QLD02/2020 (QLD02) sampled on 30/01/2020 (GISAID Accession ID; EPI_ISL_407896). This QLD02 isolate differs from the Wuhan reference strain by the following mutations: Spike P1143L, N S202N, and NS8 L84S (as classified by Pango v.4.3.1 PANGO-v1.23). These mutations are maintained in the QLD02_Δ3678_ backbone with the key difference being the targeted deletion of accessory genes ORF 3, 6, 7, and 8 as illustrated in Figure 5A. Additional strains included II) the Delta variant (B.1.617.2), specifically hCoV-19/Australia/QLD1893C/2021 collected on 05/04/2021 (GISAID accession ID EPI_ISL_2433928); III) the Omicron BA.1, named hCoV-19/Australia/QLD2639C/2021 collected on 14/12/2021 (GISAID accession ID EPI_ISL_8186598); IV) the Omicron BA.2 variant named as hCoV-19/Australia/QLD-QIMR02/2022, collected on 12/05/2022 (GISAID accession ID: EPI_ISL_15671877.2); and V) an Omicron BA.5 sub lineage BE.1 variant named hCoV-19/Australia/QLD-QIMR03/2022 (GISAID accession ID: EPI_ISL_15671874.2) collected 19/07/2022. All working virus stocks were propagated and titred on VE6-hTMPRSS2 cells using an immuno-plaque assay^29^.

### SARS CoV-2 HexaPro Spike protein expression

SARS-CoV-2 HexaPro S proteins for the Ancestral (QLD02), Delta, BA.1 and BA.2 variants were produced with the six proline substitutions and GSAS substitution of the furin cleavage site at residues 682–685^30^. Codon-optimised gBlocks™ (IDT, Singapore) encoding the mutated HexaPro S protein sequences were subcloned into the mammalian expression vectors pNBF-Hv or pNBF-Lv in-frame with an IgK signal peptide and an affinity C-tag (EPEA) for purification. Expression of these proteins was carried out in stable Chinese Hamster Ovary (CHO) bioreactors following previously described protocols^31^. Specific mutations incorporated into the HexaPro S proteins for each variant of concern were as follows: for the Alpha variant (B.1.1.7; ΔH69, ΔV70, ΔY144, N501Y, A570D, D614G, P681H, T716I, S982A, and D1118H), Delta variant (B.1.617.2; T19R, G142D, Δ156E, Δ157F, R158G, L452R, T478K, D614G, P681R, and D950N), Omicron BA.1 (A67V, D614G, D796Y, E484A, G142D, G339D, G446S, G496S, ΔH69, H655Y, ins214EPE, K417N, L212I, L981F, ΔN211, N440K, N501Y, N679K, N764K, N856K, N969K, P681H, Q493R, Q498R, Q954H, S371L, S373P, S375F, S477N, T95I, T478K, T547K, ΔV70, ΔV143, ΔY144, ΔY145, Y505H) and Omicron BA.2 variant (A27S, D405N, D614G, D796Y, E484A, G142D, G339D, H655Y, K417N, ΔL24, N440K, N501Y, N679K, N764K, N969K, ΔP25, ΔP26, P681H, Q493R, Q498R, Q954H, R408S, S371F, S373P, S375F, S477N, T19I, T376A, T478K, V213G, Y505H).

### Serum anti-Spike IgG ELISA

Antigen-specific IgG titres were quantified using the indirect ELISA method as previously described^32,33^. Briefly, high-bind protein plates (Cat: 655061, Greiner, Bio-One, Austria) were coated with 1 µg/mL of the HexaPro S antigen in PBS overnight at 4°C. The following day, plates were blocked with 5% KPL Milk Diluent/Blocking Solution (Cat: 5140-0011; SeraCare, USA) for 30 min at 37°C. Sera samples were diluted in blocking buffer starting at a dilution of 1:10 and subsequent serial dilutions at fivefold increments. A negative control of pooled pre vaccination baseline was included with blocking buffer, while a human anti-Spike monoclonal (CR3022, 50 µg/mL)^34^. After blocking 40 µL of each serial dilution was added to the ELISA plates and incubated at 37°C for 1 hour. Plates were washed three times in PBS-Tween 20 (PBS-T, 0.1%). After primary incubation and washing, 50 µL of goat anti-Human IgG Fc HRP antibody (0.5 µg/mL, Cat: A18829, Thermo Fisher Scientific, USA) with blocking buffer was added to each well. The plates were incubated for 1 hour at 37°C. After five washes with 0.1% PBST, 50µL of 3,3’,5,5’-tetramethylbenzidine (Cat: 002023, Life Technologies, USA) substrate was added. The plates were incubated at room temperature for 5 minutes and the reaction was stopped by adding 1M sulfuric acid. Absorbance was measured at 450 nm using a Varioskan LUX Microplate reader (Thermo Fisher Scientific, USA) using the SkanIt™ Software v6.0.1 (Thermo Fisher Scientific, Waltham, MA, USA).

### SARS-CoV-2 virus microneutralization assay

For the virus neutralization assays, 3x10^5^ VE6-hTMPRSS2 cells were seeded in 96-well plates in complete puromycin-containing medium as previously described and incubated overnight at 37°C with 5% CO_2_. On the day of assay, serum samples were diluted in Gibco™ minimum essential medium (MEM) supplemented with 2% (FCS) in sterile round-bottom 96-well plates. The initial dilution was 1:10, and serial 1:2 dilutions were performed. The media control wells contained MEM+2% FCS medium only. Each contained a media-only control and a pooled baseline sample from non-infected and non-vaccinated individuals from the Mater cohort. SARS-CoV-2 viruses were diluted in MEM+2% FCS and added to each well of the 96-well plates containing sera except control (virus-negative) wells. The media control wells contained MEM+2% FCS only. The plates were then incubated at room temperature for 1 hour. After incubation, 120µL of virus plus sera was transferred to VE6-TMPRSS2 cells and incubated at 37°C for 1 hour. Following the incubation period, the medium was aspirated, and 200 µL of MEM+2% FCS was added to each well. The plates were then incubated for 48 hours at 37°C. After 48 hours, the medium was aspirated, and the plates were fixed in ice-cold 80% acetone in PBS and placed at -20°C for 30 minutes before removal from the facility and processing in a PC2 laboratory. After acetone removal from the plates, the plates are dried, and PBS is added to flush out the remaining acetone. The plates were then blocked in 5% KPL solution diluted in PBS-T for 1 hour at 37°C. For primary incubation, the human Fc-linked anti-SARS-CoV-2 nucleocapsid nanobody C2^35^ was added at 1 μg/mL and incubated for 1 hour at 37°C. Plates were then washed three times with PBS-T. Subsequently, 0.2 μg/mL anti-human IgG Fc Highly Cross-Adsorbed Secondary Antibody (Cat: A18829, Thermo Fisher Scientific, USA) for 1 hour at 37°C. The plates were incubated with TMB as previously described, and the reaction was stopped by adding 1M sulfuric acid. Absorbance was measured at 450 nm using a Varioskan LUX Microplate reader (Thermo Fisher Scientific, USA).

### Generation of SARS-CoV-2 QLD02_ΔORF3678_ with Circular Polymerase Extension Reaction

Design of the attenuated recombinant SARS-CoV-2 QLD02_ΔORF3678_ virus followed that of a live-attenuated SARS-CoV-2 vaccine candidate with accessory protein deletions^36^ with some minor modifications. The virus was generated using the Circular Polymerase Extension Reaction (CPER) as per our previously described method^27^ with the following modifications (see Figure 4A). The CPER amplicon design was refined to enable seamless exchange of the complete spike gene as fragment 5, involving updated primer sets allowing for longer sequence overlaps in amplicons used for CPER assembly (Table 2).

**Table 1:** Demographic information of the Mater and UQ cohorts.

| Cohort | Group | n | Age ($\bar{x} \pm SD$) | Male | Female |
| --- | --- | --- | --- | --- | --- |
| Mater | AZ<sub>1</sub>AZ<sub>2</sub> | 62 | 43.9±12.0 | 12 (16.7%) | 50 (83.3%) |
|  | AZ<sub>1</sub>AZ<sub>2</sub>PZ<sub>3</sub> | 44 | 47.4±11.0 | 9 (20.5%) | 35 (79.5%) |
|  | AZ<sub>1</sub>AZ<sub>2</sub>PZ<sub>3</sub>PZ<sub>4</sub> | 34 | 49.8±12.5 | 7 (20.6%) | 27 (79.4%) |
|  | COVID+ | 11 | 40.8±12.6 | 4 (36.3%) | 7 (63.6%) |
| UQ | PZ<sub>1</sub>PZ<sub>2</sub> | 15 | 31.3±9.9 | 8 (53.3%) | 7 (46.7%) |
|  | PZ<sub>1</sub>PZ<sub>2</sub>PZ<sub>3</sub> | 10 | 36.2±11.2 | 6 (60.0%) | 4 (40.0%) |
|  | COVID+ | 5 | 24.1±3.2 | 4 (80.0%) | 2 (20.0%) |

**Table 2:**
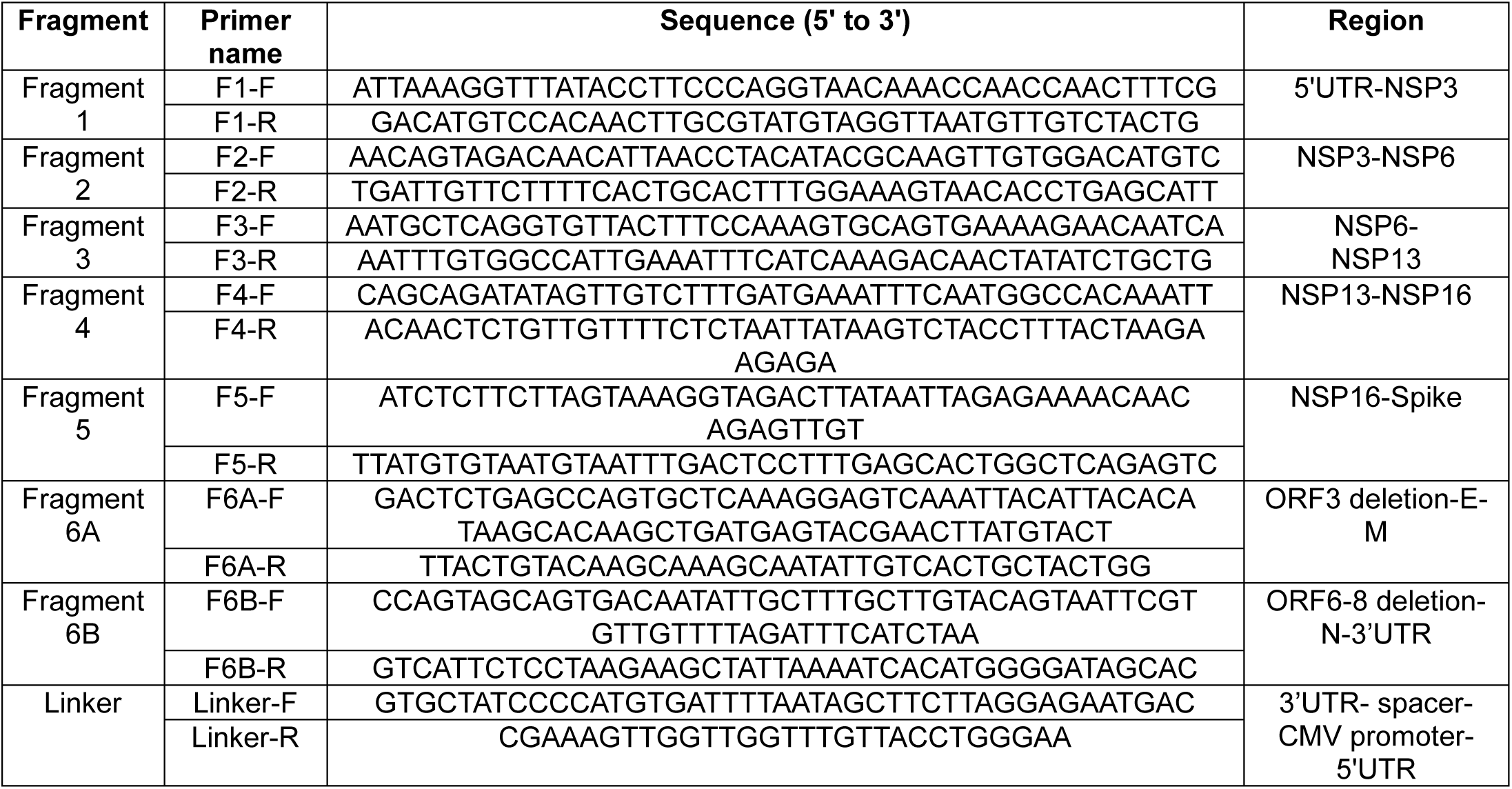
Primers used for CPER assembly of QLD02_Δ3678_ virus.

Each purified SARS-CoV-2 and linker fragments were pooled in equimolar amounts (0.1 pmol) in a 50µL CPER reaction with 2 µL of PrimeSTAR GXL DNA Polymerase (Takara Bio, Japan). The cycling conditions included: initial denaturation at 98°C for 1 minute, followed by 12 cycles of denaturation at 98°C for 10 seconds, annealing at 60°C for 20 seconds, and extension at 68°C for 10 minutes, followed by a final extension at 68°C for 10 minutes. Post-CPER assembly, the 50µL reaction mixture was transfected into HEK293T-ACE2-TMPRSS2 cells with Lipofectamine LTX Plus reagent (Invitrogen, USA) as per manufacturer’s protocol. Twelve hours post-transfection, cells were trypsinised and overlayed onto VE6-hTMPRSS2 cells. The cells were monitored daily for cytopathic effect (CPE), and culture supernatant was tested using Rapid Antigen Tests (Medomics, China) until a positive result was obtained. The recovered virus was titred on VE6-hTMPRSS2 cells using an immuno-plaque assay^29^.

### Statistical analysis

Statistical analysis and visualisation were conducted with GraphPad Prism 10.0.2 (GraphPad Software Inc., San Diego, CA, USA). The IC_50_ value of sera was calculated by regression analysis (Sigmoidal, 4PL). The IC_50_ was used to indicate the IgG antibody titer. The neutralization index was calculated by the following equation: neutralization level = 1-[(|OD_s_-OD_b_ |-OD_m_)/OD_v_-OD_m_]%. (OD_s_: absorbance of sample sera, OD_b_: absorbance of baseline sera, OD_m_: absorbance of media control, OD_v_: absorbance of virus control). Nonlinear regression (log_inhibitor_ vs. normalized response) was used to calculate the IC_50_ for neutralization assay. A linear regression analysis was undertaken to compare recombinant QLD02ΔORF3678 and QLD02 WT virus for viral neutralisation assays. For comparisons between VOCs in immunogenicity and neutralisation data, groups were first tested for normal distribution using the Kolmogorov-Smirnov test and then statistical significance between groups was determined using the Kruskal-Wallis test with Dunn’s multiple comparisons test against QLD02 (Figure 2 and 3) and against all groups (Figure 4).

**Figure 1:**
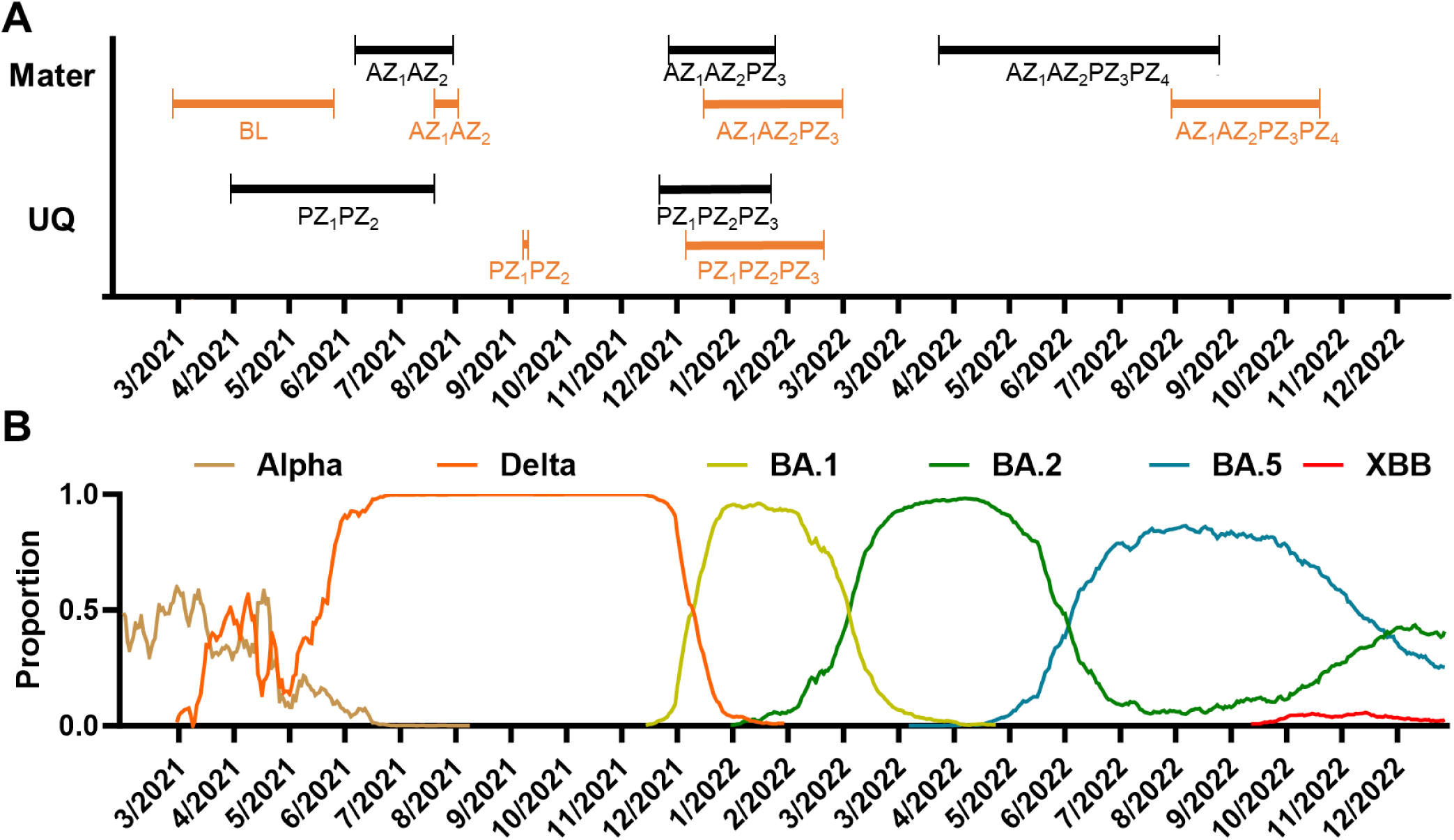
Timescale of vaccination and sera collection for study cohorts and genetic epidemiology of SARS-CoV-2 VOC lineages in Australia. A) The vaccination (black) and sera collection (orange) for AZ and PZ doses from the Mater and UQ cohorts with baseline (BL) collection indicated B) Genetic epidemiology of Australian SARS-CoV-2 variants of concern from March 2021-December 2022 generated using CoV-Spectrum^37^

**Figure 2:**
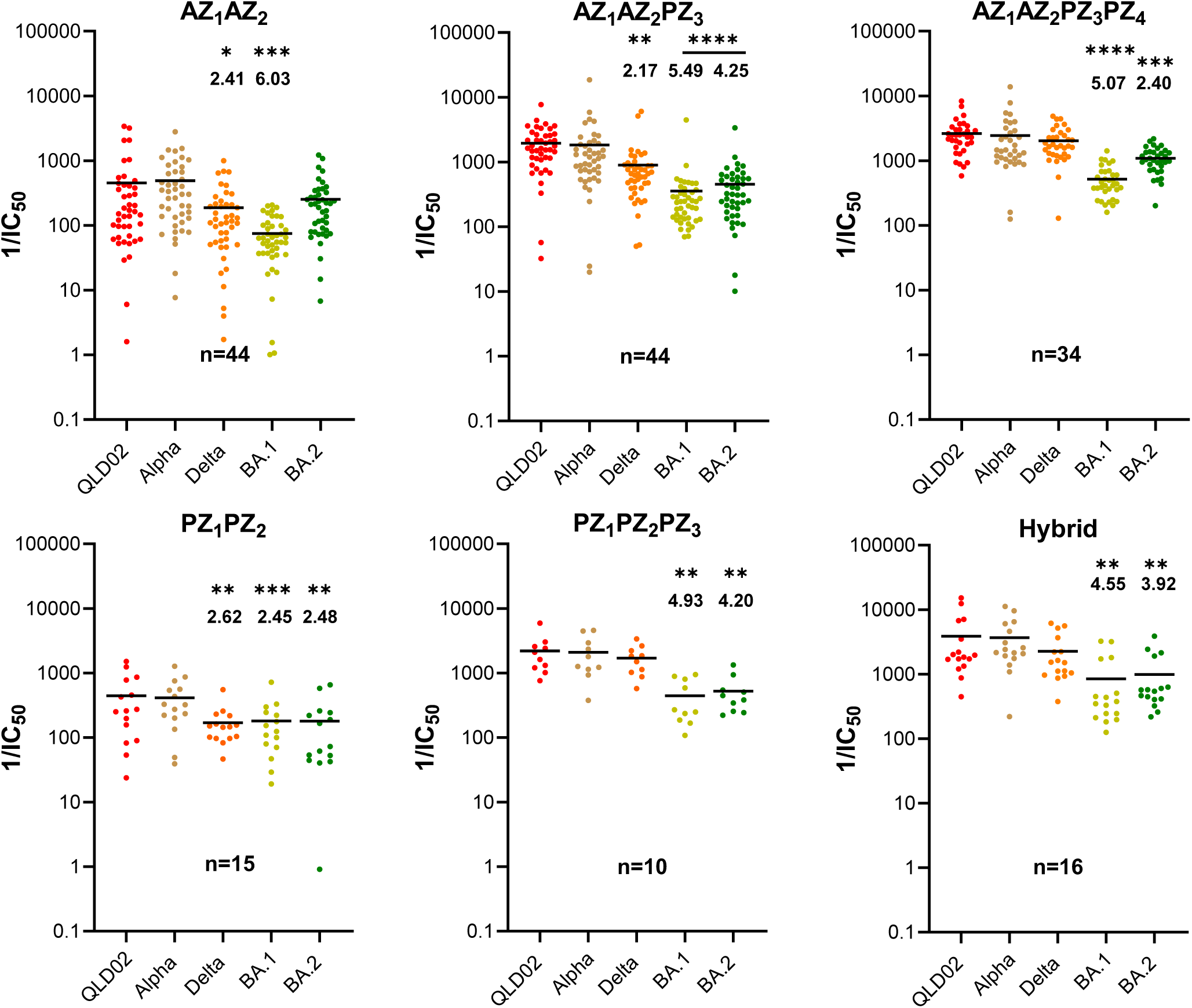
IgG antibody titres against purified Spike proteins from different SARS-CoV-2 VOCs. Average serum IgG antibody titres against HexaPro S protein derived from QLD02, Delta, and Omicron BA.1 and BA.2 variants measured by ELISA and shown by IC_50_ value. The decreased fold change compared with the ancestral virus QLD02 are shown in the figure. Statistical significance was determined using the Kruskal-Wallis test with Dunn’s multiple comparisons test against QLD02 and the fold change against QLD02 IC_50_ value is shown. Number of participants in each group is given in Table 1. (*) p < 0.05, (**) p < 0.01, (***) p < 0.001, (****) p < 0.0001 and non-significant (ns).

**Figure 3:**
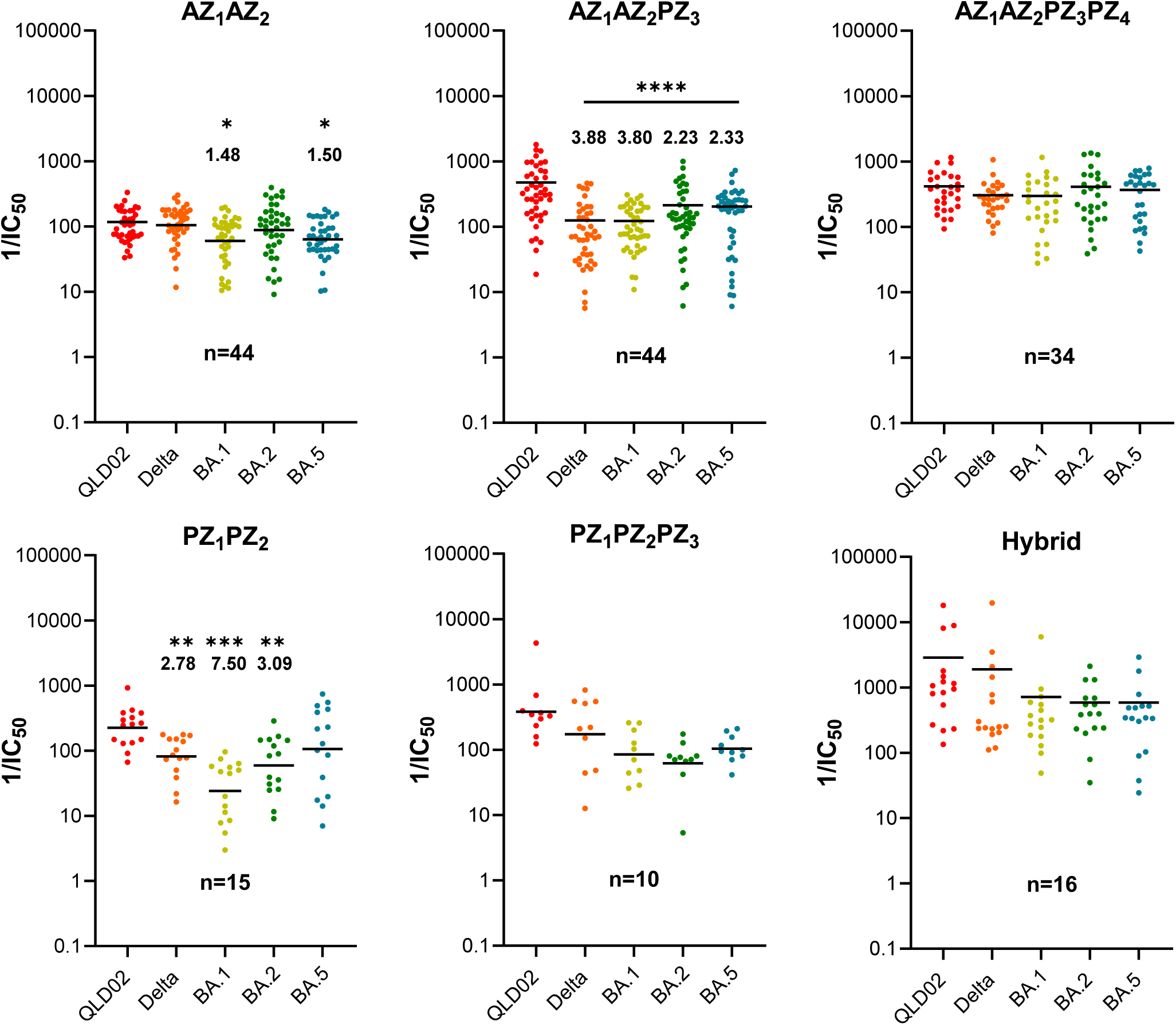
Virus-neutralizing antibody titres against SARS-CoV-2 VOCs. Sera neutralization antibody titres against QLD02, Delta, BA.1, BA.2, and BA.5 variants detected by microneutralization assay are shown by 1/IC_50_ values. Fold change reductions against VOCs compared with QLD02 ancestral virus are indicated. The mean and SEM are shown by the black bar. Statistical significance was determined using the Kruskal-Wallis test with Dunn’s multiple comparisons against QLD02. (*) p < 0.05, (**) p < 0.01, (***) p < 0.001, (****) p < 0.0001 and non-significant (ns). Number of participants in each group is given in Table 1.

**Figure 4:**
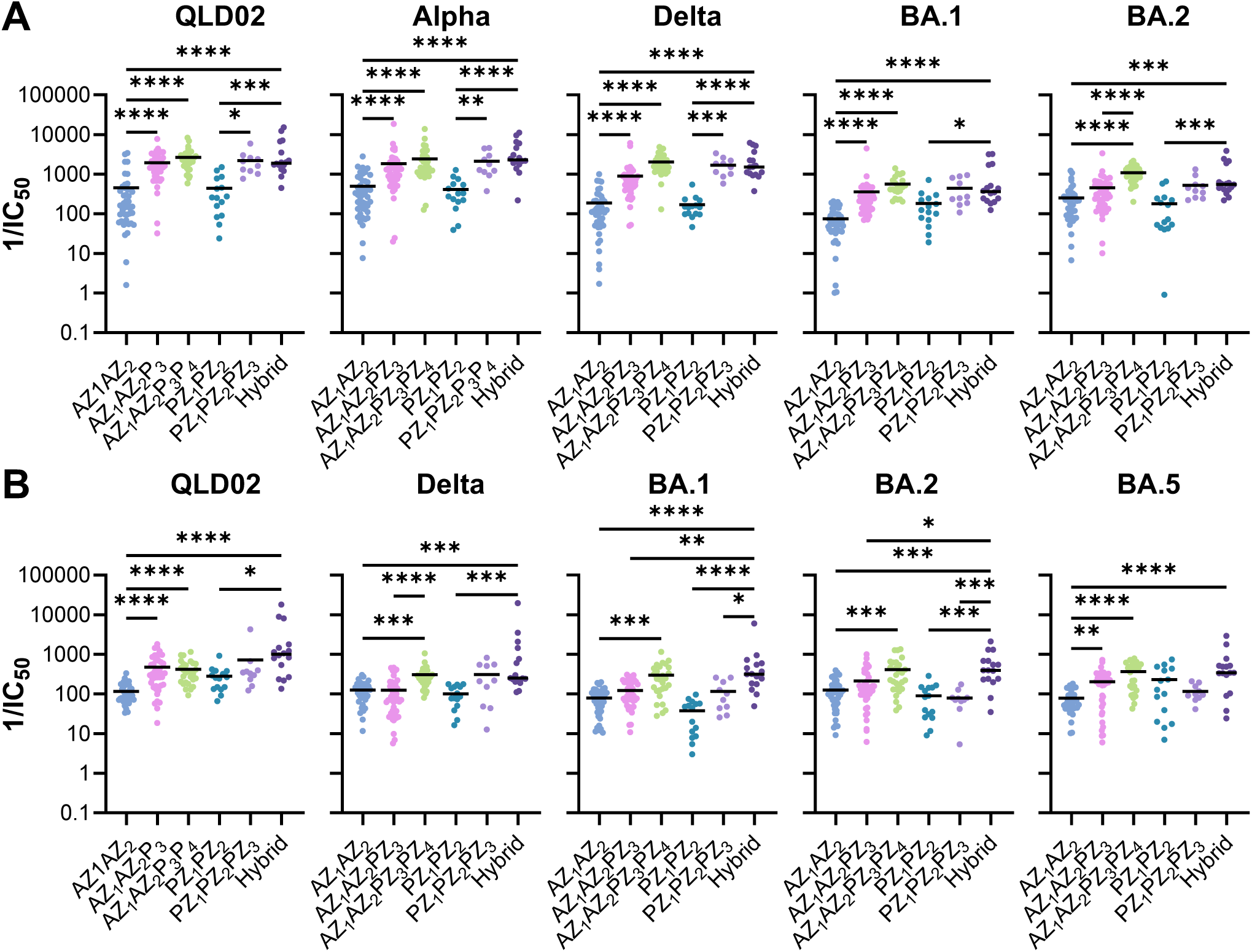
Comparison of antibody responses across different vaccination regimens against SARS-CoV-2 variants. A) Serum IgG antibody titers against HexaPro S protein derived from QLD02, Alpha, Delta, and Omicron BA.1 and BA.2 variants measured by ELISA, shown by 1/IC₅₀ values. B) Virus-neutralizing antibody titers against QLD02, Delta, and Omicron BA.1, BA.2, and BA.5 variants detected by microneutralization assay, shown by 1/IC₅₀ values. Each dot represents an individual sample, with black bars indicating mean values. Statistical significance was determined using the Kruskal-Wallis test with Dunn’s multiple comparisons for evaluating differences between vaccine regimens for each variant. (*) p < 0.05, (**) p < 0.01, (***) p < 0.001, (****) p < 0.0001 and non-significant (ns). Number of participants in each group as per groups in Figures 2 and 3.

## Results

### Sample collection and demographic characteristics of Mater Hospital and University of Queensland cohorts

The selection of healthcare workers from Mater Research Hospital and members from the University of Queensland provided complementary demographics for our study. Healthcare workers represent a priority vaccination group with higher risk of exposure to SARS-CoV-2, while the university cohort offered a broader age distribution including younger adults. This sampling strategy allowed us to evaluate vaccine responses across diverse demographics while utilizing established research networks for efficient recruitment and longitudinal follow-up. To assess the immune response elicited by vaccines or a hybrid immunity, blood samples were collected approximately six weeks after the double primary vaccinations AZ_1_AZ_2_ or PZ_1_PZ_2_ and heterologous booster vaccinations with PZ; AZ_1_AZ_2_PZ_3_ and AZ_1_AZ_2_PZ_3_PZ_4_ (Figure 1A). We conducted longitudinal sampling for sera of individuals at two locations: Mater Hospital, Brisbane, Queensland, Australia (Mater) and the University of Queensland (UQ), Brisbane, Australia, from March 2021 to December 2022 (Figure 1A). Primary vaccinations of AZ/PZ coincided with the emergence and dominance of the Delta VOC (June 2021 – December 2021) (Figure 1B). Heterologous and homologous PZ for booster vaccination, coinciding with the dominance of the Omicron BA.1 lineage (January – February 2022). Subsequent Omicron sub-lineages BA.2 and BA.5 coincided with the double booster AZ_1_AZ_2_PZ_1_PZ_2_ cohort.

At Mater, 286 healthcare workers were initially enrolled in the study, and baseline pre-vaccination sera were sampled for 191 participants. Out of these participants, sera samples were collected from 62 individuals who received double doses of the AZ vaccine (AZ_1_AZ_2_ group). Out of this group, 43 individuals received a heterologous booster of the PZ vaccine (AZ_1_AZ_2_PZ_3_), and within this subgroup, 34 individuals received a second PZ booster (AZ_1_AZ_2_PZ_3_PZ_4_). Longitudinal samples from these 34 participants were used to assess the IgG and virus-neutralising antibody responses. Additionally, 11 subjects from the AZ_1_AZ_2_PZ_3_ booster group were diagnosed with the SARS-CoV-2 virus, with sera sampled at the point of care; this group is considered a hybrid immunity group and was sampled between January to August 2022, during the Omicron BA.1 to BA.2 period, which are implied as the likely infection.

The overall Mater cohort was predominantly female (79.6%), with a mean age of 46.6 years (SD= 12.4, range 22-74 years). There were minimal differences in the average ages between the vaccination groups, with the AZ_1_AZ_2_ group being, on average, the youngest (43.9 ± 12.0 years), followed by the AZ_1_AZ_2_PZ_3_ group (47.4 ± 11.0 years) and the four-dose group (49.8 ± 11.0 years). The hybrid immunity group was 40.8 ±12.6 years.

For the PZ cohort, sera were also collected from 15 researchers at the University of Queensland (UQ). Blood draws occurred seven to nine weeks after the initial two-dose vaccination (PZ_1_PZ_2_ group) for these 15 individuals. Samples from a three-dose group that received a homologous PZ booster (PZ_1_PZ_2_PZ_3_ group, n=10 were collected two to three weeks after the third vaccine dose. Within the UQ cohort, 5 individuals (22.7%) were diagnosed with COVID-19 through positive rapid antigen or PCR tests. Of the 5 participants in the COVID+ group, two contracted SARS-CoV-2 after receiving two PZ doses, and three after receiving three PZ doses - all of whom have been allocated into the hybrid immunity group, which was collected between January – Feb 2022 and are implied as being infected with BA.1 (Figure 1B). These were pooled with COVID+ samples from the Mater cohort into a hybrid immunity group (n=16), none of these individuals had COVID prior to vaccination.

The mean age of the total UQ cohort was 30.5 ± 9.3 years, with 54.5% of participants being male (Table 1). The oldest average age was in the PZ_1_PZ_2_PZ_3_ group (36.2 ± 11.2 years), while the COVID+ group had the youngest mean age (24.1 ± 3.2 years) (Table 1).

### IgG antibody responses after vaccinations and infection

We first assessed the sera samples from individuals vaccinated with AZ and PZ vaccines and those with hybrid immunity against the purified Spike proteins of various SARS-CoV-2 variants of concern (VOCs). Serum IgG antibody titres against the HexaPro S protein derived from the ancestral QLD02 strain and Alpha, Delta, Omicron BA.1 and BA.2 variants of concern were measured using indirect ELISA. Antibody titres were represented by quantifying the half-maximal inhibitory concentration (IC_50_) values, with decreases in fold change relative to the ancestral QLD02 strain (Figure 2).

Generally, the third vaccination with the PZ vaccine elicited higher IgG antibody titres than the first two vaccinations, regardless of whether the primary regimen was AZ or PZ (Figure 2). The fourth vaccination in the cohort that received two AZ shots and one PZ shot before the fourth PZ booster did not elicit a significant increase in IgG antibody titres against the Spike proteins from ancestral virus and alpha variant while slightly improving IgG antibody titres against later VOCs (Figure 2).

Sera from vaccinated individuals, irrespective of the vaccine received, showed diminished IgG titres against Spike from Delta and Omicron VOCs, particularly against the Omicron BA.1 and BA.2 variants (Figure 2).

All vaccine groups demonstrated a consistent order of immunogenicity from highest to lowest against the Spike proteins from ancestral QLD02 to Delta, BA.2, and then BA.1 VOCs, irrespective of vaccine status. In the AZ_1_AZ_2_ group, IgG titres against Delta and Omicron BA.1 exhibited decreases by 2.41- and 6.03-fold, respectively, compared to QLD02 while decrease against BA.2 was not significant. Within the AZ_1_AZ_2_PZ_3_ cohort, the observed fold decreases were 2.17 for Delta, 5.49 for BA.1, and 4.25 for BA.2. Participants in the AZ_1_AZ_2_PZ_3_PZ_4_ group showed less of a decrease in titres against VOCs, with a 5.07 for BA.1, and 2.40 for BA.2. The hybrid immunity group exhibited a 4.55-fold decrease for BA.1 and 3.92-fold decrease for BA.2 while no significant decrease was detected for Delta variant.

For the PZ-vaccinated cohorts, the PZ_1_PZ_2_ group demonstrated a 2.62-fold decrease in IgG titres against Delta, 2.45 for BA.1, and 2.48 for BA.2 compared to QLD02. The PZ_1_PZ_2_PZ_3_ group had a 4.93-fold decrease for BA.1 and 4.20 for BA.2, while no significant difference in antibody titres were detected against the Delta variant.

### Neutralising antibody responses against SARS-CoV-2 VOCs

The virus neutralizing capacity of sera was tested against a panel of SARS-CoV-2 strains, including the ancestral strain QLD02 and the variants of concern (VOCs) Delta, and the Omicron lineages BA.1, BA.2, and BA.5, across different vaccination regimens: AZ_1_AZ_2_, AZ_1_AZ_2_PZ_3_, AZ_1_AZ_2_PZ_3_PZ_4_, PZ_1_PZ_2_, PZ_1_PZ_2_PZ_3_ and the hybrid immunity group (Figure 3). The Alpha VOC was not tested for neutralization as it did not show significant differences to QLD02 in immunogenicity tested by ELISA in these sera samples (Figure 2). BA.5 VOC was added to the testing as the latest Omicron variant circulating at the time of the study. The neutralizing antibody titres were quantified using a microneutralization assay, expressed as 1/IC_50_ values, representing the serum dilution required to inhibit 50% of the virus infectivity.

In the AZ_1_AZ_2_ group, the sera showed a slightly reduced but significant neutralization against BA.1 and BA.5 with fold changes of 1.48 and 1.50, respectively. In contrast, the AZ_1_AZ_2_PZ_3_ group exhibited a significant reduction in neutralization capacity against all VOCs ranging from 2.23 (BA.2) to 3.88 (Delta). Notably, the AZ_1_AZ_2_PZ_3_PZ_4_ group demonstrated no significant virus neutralization differences against all tested VOCs, including all Omicron variants, clearly demonstrating the benefit from an additional PZ booster.

Participants in the PZ_1_PZ_2_ group displayed the greatest reduction in neutralizing titres against BA.1 (7.5-fold, p < 0.001), followed by BA.2 (3.09-fold, p < 0.01) and Delta (2.78-fold, p < 0.01), with no significant reduction in neutralization for the BA.5 variant (Figure 3). The PZ_1_PZ_2_PZ_3_ cohort showed no significant difference in neutralization for all VOCs, although this could be due to a smaller number of samples (Figure 3).

For the hybrid immunity group, no significant differences were also observed for all VOCs despite visual difference, although this could also be due to the smaller cohort size (Figure 3).

Overall, the results indicate that while initial vaccinations with AZ or PZ vaccines showed a tendency for decreased neutralization of Delta and Omicron OCs, additional booster(s) with PZ vaccine appear to elicit more effective neutralization responses against theses VOCs.

### Comparisons of antibody responses for each virus variant across different vaccination regimens

To understand whether different vaccination regimens elicited varying levels of antibody responses, we compared IgG and virus-neutralizing antibody titers across all vaccination groups for each SARS-CoV-2 variant (Figure 4). When comparing IgG antibody responses against the ancestral QLD02 strain (Figure 4A), we observed a general trend toward higher antibody titers in groups that received three or four vaccine doses compared to those who received two doses, regardless of vaccine type (AZ or PZ). Similar patterns were observed for Alpha, Delta, and Omicron variants, although the differences between vaccination regimens were less pronounced for Omicron BA.1 and BA.2.

Interestingly, when comparing the two double-dose regimens (AZ₁AZ₂ vs. PZ₁PZ₂), no statistically significant differences in antibody titers were observed against any of the tested variants, suggesting comparable immunogenicity between the two vaccine platforms when administered as a primary series. Similarly, heterologous boosting (AZ₁AZ₂PZ₃) did not show significant advantages or disadvantages compared to homologous boosting (PZ₁PZ₂PZ₃) in terms of antibody titers against any variant.

For virus neutralization (Figure 4B), the patterns were largely consistent with the IgG results. The hybrid immunity group generally displayed the highest neutralization titers across all variants, although these differences were not always statistically significant, potentially due to the smaller sample size in this group. The four-dose regimen (AZ₁AZ₂PZ₃PZ₄) also showed robust neutralization against all variants tested, including the Omicron sublineages.

It is important to note that demographic differences between the cohorts, particularly age disparities between the Mater and UQ groups, may confound these comparisons. The UQ cohort (predominantly receiving PZ vaccines) had a younger mean age (30.5 ± 9.3 years) compared to the Mater cohort (predominantly receiving AZ vaccines initially, with mean age 46.6 ± 12.4 years), which might influence immune responses independent of vaccination regimen.

### Generation of an attenuated SARS-CoV-2 virus with deletion of all accessory genes and its application for virus-neutralization assays

Currently, live virus neutralisation assays of sera samples need to be undertaken in facilities with BSL3/PC3 containment. SARS-CoV-2 virus with deletion of all accessory genes, ORF 3, 6, 7 and 8 (ΔORF3678), was recently developed and was shown to be severely attenuated in interferon response-competent human cells and in animals while still allowing efficient propagation in interferon-deficient Vero E6 cells^36^. Such attenuated viruses allow neutralization assays to be performed in less stringent BSL2/PC2 containment facilities while retaining the characteristic features of SARS-CoV-2 infection^38^.

Herein, we employed our previously developed for SARS-CoV-2 circular polymerase extension reaction (CPER)^27^ to generate SARS-CoV-2 virus with deletion of ORFs 3, 6, 7 and 8 based on ancestral QLD02 strain QLD02_ΔORF3678_. To facilitate rapid adaptation of neutralization analysis with future VOCs we reconfigured our previous CPER amplicon strategy to allow seamless swapping of the complete Spike genes from any future VOCs in fragment 5 (Figure 5A). Using our new CPER design, we recovered the QLD02_ΔORF3678_ virus following transfection of CPER assembly mix into HEK293-ACE2-TMPRSS2 cells which were co-cultured with VE6-hTMPRSS2 cells to recover and amplify the virus yielding viral titre of 1.34 x 10^6^ FFU/mL. RT-PCR from isolated viral RNA confirmed ORF3678 deletion (Figure 5B). Immunoplaque assay showed that the virus produced smaller foci compared to the parental QLD02 virus in Vero E6 cells (Figure 5C).

**Figure 5:**
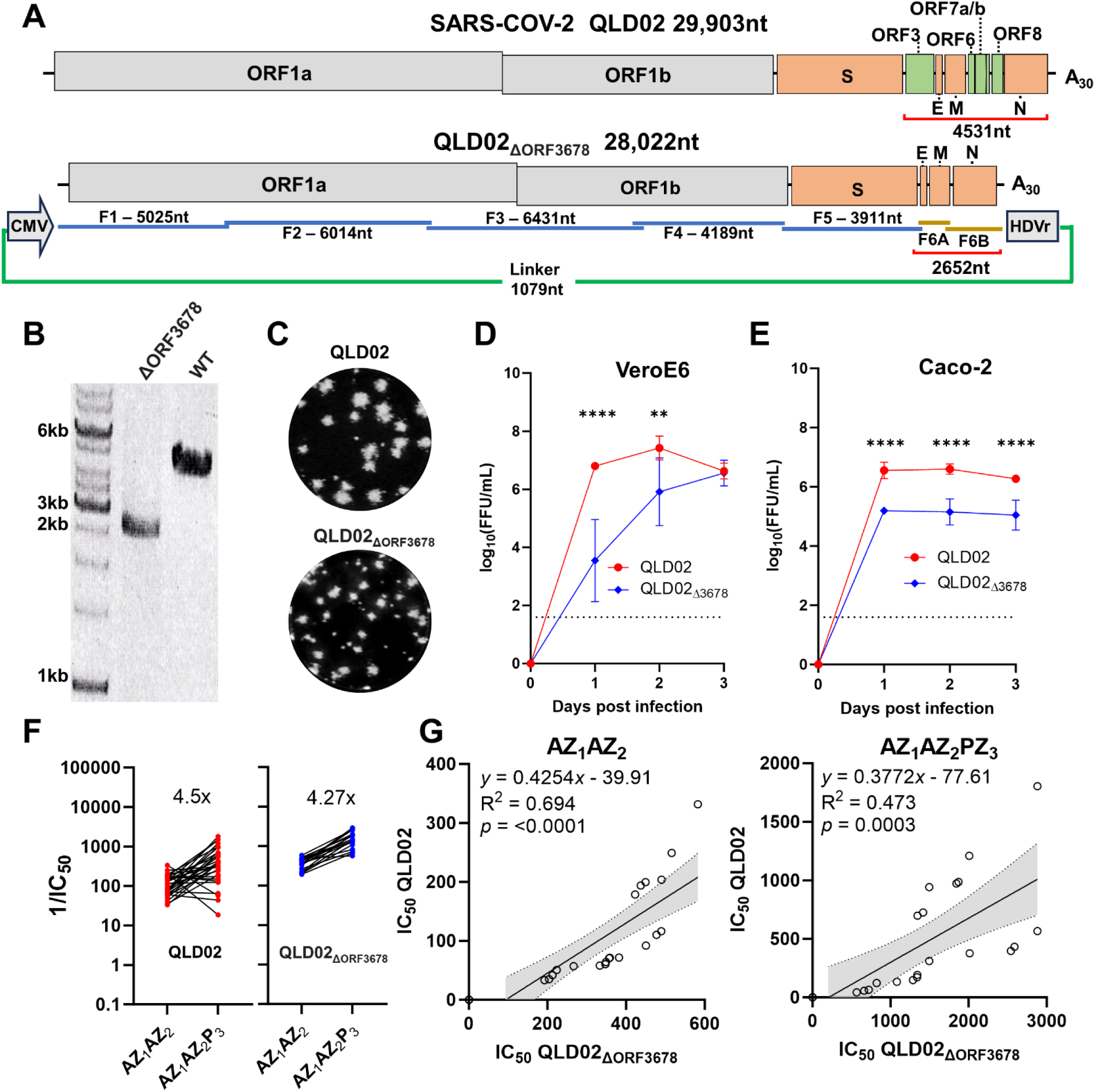
Generation and characterisation of an attenuated recombinant QLD02_ΔORF3678_ virus as a substitute for native SARS-CoV-2 in virus-neutralization assay. A) Schematics of the SARS-CoV-2 QLD02 and the recombinant QLD02_ΔORF3678_ virus genomes and amplicon fragments used for CPER assembly. Red lines with nucleotide sizes show RT-PCR amplicons of fragment 6 and the linker construct (green line) for the QLD02 and QLD02_ΔORF3678_ viruses shown in B). Linker fragment is shown not to scale and contains the Cytomegalovirus (CMV) promoter and Hepatitis delta virus ribozyme (HDVr). C) Immuno-plaque assay with anti-SARS-CoV-2 nucleocapsid nanobody at 20 hours post-infection. Growth kinetics of WT QLD02 (red) and QLD02_ΔORF3678_ (blue) viruses over a 3-day time course in D) Vero E6 cells and E) Caco2 cells infected at a multiplicity of infection MOI = 0.01, n = 2 independent experiments with three replicates in each, statistical analysis was performed by two-way analysis of variance with Tukey’s multiple comparisons test against WT virus. Mean values for each virus at each time point are shown ± SD. F) Sera neutralization antibody titres against QLD02 and QLD02_ΔORF3678_ SARS-CoV-2 virus by microneutralization assay, shown as fold reduction in viral load compared to control as IC_50_ values, for 20 matched samples from the AZ₁AZ₂ and AZ₁AZ₂PZ₃ groups. G) The linear relationship between matched samples of IC50 values from both AZ₁AZ₂ and AZ₁AZ₂PZ₃ samples (n=20) and baseline unvaccinated samples (n=3). The linear regression function, coefficient of determination (R²), and p-value for the function are given, with 95% confidence bands for the line of best fit.

We then compared the growth of WT QLD02 and QLD02_ΔORF3678_ viruses in interferon-deficient Vero E6 cells and interferon-competent human Caco-2 cells at a MOI 0.01 over a three-day time course. Both viruses showed pronounced growth in Vero E6 cells (Figure 5D), with the QLD02_ΔORF3678_ exhibiting delayed replication and significant differences between titres of the two viruses in the first two days (Day 1 *p*<0.0001, Day 2 *p*=0.0020, two-way ordinary ANOVA with Tukey’s multiple comparisons test) while reaching comparable titres by day 3 (*p*=0.9997) (Figure 5D). In the interferon competent human Caco2 cell line, QLD02_ΔORF3678_ grew to significantly lower viral titres than WT QLD02 at all time points (Figure 5E).

To examine if QLD02_ΔORF3678_ virus could be used as a surrogate for virus-microneutralization assays, sera samples from 20 individuals that matched between AZ_1_AZ_2_ and AZ_1_AZ_2_PZ_3_ groups were examined for virus neutralisation assays with the ΔORF3678 virus and compared with neutralization by WT QLD02 virus. The serum neutralization titres for the ΔORF3678 compared to the WT QLD02 virus were on average 3.29-fold higher for the AZ_1_AZ_2_ samples and 3.11-fold higher for the AZ_1_AZ_2_PZ_3_ samples indicating higher assay sensitivity. In addition, the assay with QLD02_ΔORF3678_ virus showed lower variability between individual samples than WT QLD02 virus while maintaining similarly increased neutralization between pre- and pos-booster immunization groups (Figure 5F).

To further examine the harmony between individual samples being assayed for IC_50_ using both viruses we plotted the relationship between IC_50_ values of the WT virus and ΔORF3678 virus for the same samples (Figure 5G). Three pre-immunisation sera samples were added to calibrate the baseline at IC_50_ 0. Generally, there was a strong linear relationship between WT QLD02 IC_50_ values and the ΔORF3678 virus for the AZ_1_AZ_2_ (R^2^ = 0.694) and a slightly weaker relationship for the AZ_1_AZ_2_PZ_3_ samples (R^2^ = 0.473). There was also a significant linear relationship between IC_50_ values for the WT virus and ΔORF3678 virus for both pre- and post-booster immunization groups as indicated by the slope test (AZ_1_AZ_2,_ p = <0.0001; AZ_1_AZ_2_PZ_3_, p = 0.0003). The data therefore show that using the ΔORF3678 virus in virus neutralisation assays recapitulates results generated using WT QLD02 virus, and therefore an attenuated ΔORF3678 virus could be used as a suitable substitute for the WT virus in neutralization assays in BSL-2/PC2 facilities.

## Discussion

Here we examined the total IgG and virus-neutralizing antibodies in the longitudinal sera from participants undergoing SARS-CoV-2 vaccinations with Vaxzevria (AstraZeneca: AZ) and Comirnaty (Pfizer: PZ) vaccines and some of them infected with SARS-CoV-2 in two cohorts in Queensland, Australia. We show that similar to other published findings, Spike-specific IgG and virus-neutralising antibody titres were reduced for VOCs Delta and Omicron sublineages. Neutralization assays with natural isolates of SARS-CoV-2 revealed reductions in neutralizing antibody titers against Omicron sublineages: in the AZ_1_AZ_2_ group with a follow up single PZ (AZ_1_AZ_2_PZ_3_) and double PZ (AZ_1_AZ_2_PZ_3_PZ_4_) boosters, titers were reduced by 3.80-fold and 1.40-fold for BA.1, respectively, and 2.23-fold and 1.02-fold for BA.2, respectively. For BA.5, the reduction was 2.33-fold in the AZ_1_AZ_2_PZ_3_ group and 1.12-fold in AZ_1_AZ_2_PZ_3_PZ_4_ group. Clearly, the second booster with PZ vaccine in these groups substantially reduced the difference in neutralization between different VOCs. The modest reductions we observed in neutralization against Delta and Omicron variants (BA.1, BA.2, and BA.5) compared to the ancestral virus are consistent with previous reports using both wild-type virus and pseudotyped virus neutralization assays^39,40^. In primary vaccination regimens, much more severe reductions were observed when pseudoviruses were used in neutralization assays. For example, neutralization of a pseudovirus with BA.1 Spike protein reduced by 33-fold for PZ_1_PZ_2_ and 14-fold for AZ_1_AZ_2_^41^. In comparison, PRNT virus neutralization using infectious virus showed 7.5-fold reduction in PZ_1_PZ_2_PZ_3_ cohort compared to ancestral virus^22^. This argues for remaining need for virus-neutralization assays to more accurately estimate antibody responses.

Our study deliberately focused on the 2021-2022 timeframe to capture immune responses temporally matched to the circulating variants experienced by our cohorts at the time. This approach maintains the critical temporal relationship between vaccination/infection and variant exposure, avoiding potential confounding factors in interpreting humoral IgG responses. The value of this design lies in matching seropositivity measurements with contemporaneous variants, providing an accurate snapshot of population immunity during this specific period.

### Antigen heterogeneity between prefusion locked and wild type vaccine Spike protein

Cryo-EM studies have indicated that the SARS-CoV-2 spike protein is exceptionally flexible and displays multiple prefusion conformations^42^, with its three RBDs assuming different positions: an “up” state that allows receptor access, and a “down” state that prevents receptor access. The Pfizer/BioNTech’s BNT162b2 messenger RNA vaccine (PZ) encodes a mutant spike protein and is formulated as an RNA–lipid nanoparticle of nucleoside-modified mRNA^3^. The Spike protein in BNT162b2 vaccine is locked in a prefusion conformation by two proline substitutions at residues 986-987^43^. The Oxford/AstraZeneca’s ChAdOx1-S non-replicating adenovirus-vectored vaccine (AZ) expresses the SARS-CoV-2 spike protein gene with no mutations^4^. The HexaPro is an engineered version of the Spike protein with six proline substitutions located at positions F817P, A892P, A899P, A942P, K986P, V987P and a mutated furin cleavage site (RRAR to GSAG). These changes have been shown to result in stable prefusion spike trimers^43,44^. It’s important to note that our IgG assays utilised plates coated with HexaPro proteins, engineered to stabilize the Spike protein in the prefusion conformation. In contrast, the AZ vaccine encodes a Spike protein without heterologous mutations that bias into the prefusion state, allowing for greater conformational flexibility. This difference may lead to weaker or altered binding of IgGs elicited by the AZ vaccine to HexaPro-coated assays, potentially influencing the observed differences in immunogenicity between vaccine groups. Future studies could investigate the ratio of prefusion to postfusion specific IgGs to provide a more wholistic understanding of the humoral immune response to different vaccine types.

### Individual mutations in Spike protein contributing to antibody escape

The spike protein of SARS-CoV-2 has evolved to maintain a delicate balance between engaging the ACE2 receptor for viral entry and mediating escape against neutralizing epitopes^14^. This dual functionality of the spike protein underscores its central role in the ongoing evolution of SARS-CoV-2 and its variants. In the spike protein of SARS-CoV-2 VOCs, several key amino acids have been identified that mediate escape in neutralization. In the BA.5 variant, the L452R mutation is associated with a reduction in neutralisation in both convalescent and immunised persons^45,46^. The E484A mutation usually results in a high degree of resistance to several neutralising monoclonal antibodies and polyclonal sera^13^. In the Delta variant B.1.617.2, the L452R and T478K mutations are associated with moderate resistance with immune escape and enhanced viral entry into cells^47^. In the Omicron BA.1 and BA.2 variants, the E484A, N501Y, K417N and Δ69-70 mutations are associated with immune escape^48,49^. The G446S mutations in BA.2 are thought to be primarily responsible for its enhanced resistance to neutralizing antibodies^50^. Both S371L/F mutations in BA.1/BA.2 are associated with immune escape^51^. The combinatorial mosaic of immune-resistant residues between these groups in the most recent VOC we used in our study (BA.5) includes E484A, N501Y, K417N, and L452R. While these changes may contribute to the antibody escape, our data show that two immunizations with AZ vaccine followed by two additional boosters with PZ vaccine elicit antibody response that is able to efficiently neutralise Omicron VOCs.

### Recombinant SARS-CoV-2 with deletion of all accessory genes as a tool allowing to conduct research and serological testing under biosafety level 2 conditions

In this study, we successfully generated a recombinant, attenuated SARS-CoV-2 virus with deletions in the accessory genes ORF 3, 6, 7, and 8 (QLD02_Δ3678_). The QLD02_Δ3678_ mutant exhibited delayed and reduced replication on days 1 and 2 in Vero E6 cells and demonstrated reduced replication across all days in the Caco2 cell line. Despite this attenuation, the QLD02_Δ3678_ SARS-CoV-2 neutralization assay closely resembles the virus neutralization assays (VNA) from wild-type SARS-CoV-2. This similarity allows the assay to generate neutralization results that closely approximate the gold standard plaque reduction neutralization test (PRNT).

Our findings indicate that the QLD02_Δ3678_ virus replicates robustly in Vero E6 cells, achieving viral titres exceeding 10^6^ FFU/mL. This high replication efficiency facilitates the large-scale production of high-quality viral stocks, ensuring a sufficient supply and reducing costs for serological testing. At the same time, demonstrated significant attenuation of such virus in human cells and in laboratory animals^36^ allows to conduct these assays under biosafety level 2 (BSL-2) conditions^38^. This presents a significant advantage, as BSL-2 facilities are more widely available and easier to work with than the higher containment BSL-3 laboratories. Additionally, almost all clinical laboratories operate at BSL-2, thus broadening the accessibility and application of this assay.

Incorporating the Δ3678 mutant virus into serological testing workflows offers several other significant advantages. Firstly, the attenuated Δ3678 SARS-CoV-2 closely resembles the authentic virus regarding structural characteristics, maturation, and infection pathways. This ensures that the neutralization results are reliable and comparable to those obtained using authentic SARS-CoV-2. It is also important to maintain the remaining viral genetic backbone, i.e. the genome outside spike, in neutralisation assays as it has been shown that, at least for BA.1, attenuating mutations outside the spike protein i.e. NSP6^52^ affect virus growth kinetics *in vitro,* which may confound neutralisation assays.

It has been shown that IgG and neutralizing antibody responses, specifically ratio of IgG1/IgG3 positively correlate with neutralising ability for SARS-CoV-2 infection ^53–55^. It has also been shown that for BA.1, there is a significant reduction in the relationship between IgG and neutralisation assays ^56^. This further highlights the need for a workable virus-neutralization test, considered as the gold standard for measuring neutralising antibodies against SARS-CoV-2 infection^57^.

The flexibility of the Δ3678 platform and the updated CPER design allows for the potential incorporation of reporter genes or scaling up to 96-well formats, which could significantly enhance the throughput of virus neutralization assays. Additionally, our updated CPER format now includes the spike gene on an individual fragment (F5), which allows for the rapid exchange of the spike gene from different VOCs. While a similar ΔORF3678 recombinant virus system has already been developed^36,38^, generation of recombinant viruses with exchanged Spike genes from VOCs using this approach requires a multistep process involving cloning and propagation of cloned fragments in bacteria, followed by *in vitro* DNA ligation and *in vitro* transcription of a large viral RNA. In contrast, our CPER approach does not require cloning, propagation in bacteria, and *in vitro* RNA transcription, thus facilitating the rapid generation of recombinant viruses containing spike genes from future VOCs.

This adaptability is particularly valuable for high-throughput screening in research and clinical settings, enabling rapid and efficient testing of large numbers of samples. In summary, developing a highly sensitive and robust live-attenuated QLD02_Δ3678_ SARS-CoV-2 assay that can be used in BSL-2 laboratories improves COVID-19 serological testing. This assay maintains high reliability and accessibility, making it an invaluable tool for ongoing and future research and diagnostic efforts against SARS-CoV-2.

While multiple studies have examined vaccine responses against SARS-CoV-2 variants, our work makes several unique contributions. Firstly, we provide valuable longitudinal data from Australian cohorts during a critical vaccination period (2021-2022), with particular emphasis on heterologous vaccination schedules (AZ followed by PZ boosters) that were common in Australia but less extensively studied globally. Secondly, our attenuated QLD02Δ3678 virus system enables neutralization studies in BSL2 facilities while maintaining concordance with wild-type virus results, thus addressing a significant practical barrier in SARS-CoV-2 research. Thirdly, our CPER-based system represents a significant methodological advancement for generating recombinant viruses with VOC Spike genes, eliminating the need for time-consuming cloning and bacterial propagation steps. Our CPER-based approach for generating QLD02Δ3678 offers advantages for incorporating Spike genes from emerging VOCs. To incorporate a new VOC spike sequence, the spike gene from the variant of interest would be amplified using primers F5-F and F5-R (Table 2), which contain overlapping regions with adjacent fragments. This amplicon would replace Fragment 5 in the CPER reaction, allowing integration of any VOC Spike gene into the QLD02 backbone with deletion of ORFs 3, 6, 7, and 8. Finally, our comprehensive comparison of immunogenicity and neutralization against multiple Omicron sublineages within the same cohorts provides important insights into variant-specific responses across different vaccination regimens.

## Acknowledgements

This study was supported by the NHMRC Ideas Grant (2012883) to AAK, AAA and LDB and Australian Infectious Diseases Research Center (AIDRC) Seed Grant to AAK. We thank Frederick Moore and Alyssa Pyke, Queensland Health, Brisbane, for providing SARS-CoV-2 virus isolates. We acknowledge the samples obtained through the David Serisier Research Biobank – Mater Research, and Megan L. Martin and Timothy Nguyen for their assistance in sample processing. We are grateful to Xuping Xie and Pei-Yong Shi for providing advice on construction of ΔORF3678 virus.

## Conflict of Interest statement

All authors declare that they have no conflicts of interest.

## Author Contributions

R.H.P., A.A.K. conceptualization; R.H.P., Y.S., J.D.J.S. investigation; C.L.D.M., N.M., A.A.A., A.I., B.L., D.A.M., D.R., A.S., D.W. resources; S.A., S.K. administration; R.H.P., A.A.K. project administration; A.A.A., L.D.B., A.A.K. funding acquisition. R.H.P., Y.S., J.D.J.S., C.L.D.M., N.M., A.A.A., A.I., B.L., D.A.M., D.R., A.S., S.A., S.K., D.W., L.D.B., and A.A.K. writing–review & editing; R.H.P., Y.S. writing–original draft.

## Notes

### Competing Interest Statement

The authors have declared no competing interest.

### Summary of Updates

Updated to include new figure 4 (comparisons across vaccine regimens), updated CPER methodology, discussion and Author Contributions Conflict of Interest statements.

## References

1. Lou, B., Li, T.D., Zheng, S.F., Su, Y.Y., Li, Z.Y., Liu, W., Yu, F., Ge, S.X., Zou, Q.D., Yuan, Q., et al. (2020). Serology characteristics of SARS-CoV-2 infection after exposure and post-symptom onset. Eur Respir J 56. 10.1183/13993003.00763-2020.

2. Carvalho, T., Krammer, F., and Iwasaki, A. (2021). The first 12 months of COVID-19: a timeline of immunological insights. Nat Rev Immunol 21, 245–256. 10.1038/s41577-021-00522-1.

3. Polack, F.P., Thomas, S.J., Kitchin, N., Absalon, J., Gurtman, A., Lockhart, S., Perez, J.L., Perez Marc, G., Moreira, E.D., Zerbini, C., et al. (2020). Safety and Efficacy of the BNT162b2 mRNA Covid-19 Vaccine. N Engl J Med 383, 2603–2615. 10.1056/NEJMoa2034577.

4. Voysey, M., Clemens, S.A.C., Madhi, S.A., Weckx, L.Y., Folegatti, P.M., Aley, P.K., Angus, B., Baillie, V.L., Barnabas, S.L., Bhorat, Q.E., et al. (2021). Safety and efficacy of the ChAdOx1 nCoV-19 vaccine (AZD1222) against SARS-CoV-2: an interim analysis of four randomised controlled trials in Brazil, South Africa, and the UK. Lancet 397, 99–111. 10.1016/S0140-6736(20)32661-1.

5. Shrotri, M., Fragaszy, E., Nguyen, V., Navaratnam, A.M.D., Geismar, C., Beale, S., Kovar, J., Byrne, T.E., Fong, W.L.E., Patel, P., et al. (2022). Spike-antibody responses to COVID-19 vaccination by demographic and clinical factors in a prospective community cohort study. Nat Commun 13, 5780. 10.1038/s41467-022-33550-z.

6. Erice, A., Varillas-Delgado, D., and Caballero, C. (2022). Decline of antibody titres 3 months after two doses of BNT162b2 in non-immunocompromised adults. Clin Microbiol Infect 28, 139 e131–139 e134. 10.1016/j.cmi.2021.08.023.

7. Ward, H., Whitaker, M., Flower, B., Tang, S.N., Atchison, C., Darzi, A., Donnelly, C.A., Cann, A., Diggle, P.J., Ashby, D., et al. (2022). Population antibody responses following COVID-19 vaccination in 212,102 individuals. Nat Commun 13, 907. 10.1038/s41467-022-28527-x.

8. Wei, J., Pouwels, K.B., Stoesser, N., Matthews, P.C., Diamond, I., Studley, R., Rourke, E., Cook, D., Bell, J.I., Newton, J.N., et al. (2022). Antibody responses and correlates of protection in the general population after two doses of the ChAdOx1 or BNT162b2 vaccines. Nat Med 28, 1072–1082. 10.1038/s41591-022-01721-6.

9. Seow, J., Graham, C., Merrick, B., Acors, S., Pickering, S., Steel, K.J.A., Hemmings, O., O’Byrne, A., Kouphou, N., Galao, R.P., et al. (2020). Longitudinal observation and decline of neutralizing antibody responses in the three months following SARS-CoV-2 infection in humans. Nat Microbiol 5, 1598–1607. 10.1038/s41564-020-00813-8.

10. Toh, Z.Q., Anderson, J., Mazarakis, N., Neeland, M., Higgins, R.A., Rautenbacher, K., Dohle, K., Nguyen, J., Overmars, I., Donato, C., et al. (2022). Comparison of Seroconversion in Children and Adults With Mild COVID-19. JAMA Netw Open 5, e221313. 10.1001/jamanetworkopen.2022.1313.

11. Tong, M.Z., Sng, J.D., Carney, M., Cooper, L., Brown, S., Lineburg, K.E., Chew, K.Y., Collins, N., Ignacio, K., Airey, M., et al. (2023). Elevated BMI reduces the humoral response to SARS-CoV-2 infection. Clin Transl Immunology 12, e1476. 10.1002/cti2.1476.

12. Menni, C., May, A., Polidori, L., Louca, P., Wolf, J., Capdevila, J., Hu, C., Ourselin, S., Steves, C.J., Valdes, A.M., and Spector, T.D. (2022). COVID-19 vaccine waning and effectiveness and side-effects of boosters: a prospective community study from the ZOE COVID Study. Lancet Infect Dis 22, 1002–1010. 10.1016/S1473-3099(22)00146-3.

13. Liu, Z., VanBlargan, L.A., Bloyet, L.M., Rothlauf, P.W., Chen, R.E., Stumpf, S., Zhao, H., Errico, J.M., Theel, E.S., Liebeskind, M.J., et al. (2021). Identification of SARS-CoV-2 spike mutations that attenuate monoclonal and serum antibody neutralization. Cell Host Microbe 29, 477–488 e474. 10.1016/j.chom.2021.01.014.

14. Harvey, W.T., Carabelli, A.M., Jackson, B., Gupta, R.K., Thomson, E.C., Harrison, E.M., Ludden, C., Reeve, R., Rambaut, A., Consortium, C.-G.U., et al. (2021). SARS-CoV-2 variants, spike mutations and immune escape. Nat Rev Microbiol 19, 409–424. 10.1038/s41579-021-00573-0.

15. Volz, E., Mishra, S., Chand, M., Barrett, J.C., Johnson, R., Geidelberg, L., Hinsley, W.R., Laydon, D.J., Dabrera, G., O’Toole, A., et al. (2021). Assessing transmissibility of SARS-CoV-2 lineage B.1.1.7 in England. Nature 593, 266–269. 10.1038/s41586-021-03470-x.

16. Meng, B., Kemp, S.A., Papa, G., Datir, R., Ferreira, I., Marelli, S., Harvey, W.T., Lytras, S., Mohamed, A., Gallo, G., et al. (2021). Recurrent emergence of SARS-CoV-2 spike deletion H69/V70 and its role in the Alpha variant B.1.1.7. Cell Rep 35, 109292. 10.1016/j.celrep.2021.109292.

17. Korber, B., Fischer, W.M., Gnanakaran, S., Yoon, H., Theiler, J., Abfalterer, W., Hengartner, N., Giorgi, E.E., Bhattacharya, T., Foley, B., et al. (2020). Tracking Changes in SARS-CoV-2 Spike: Evidence that D614G Increases Infectivity of the COVID-19 Virus. Cell 182, 812–827 e819. 10.1016/j.cell.2020.06.043.

18. Mlcochova, P., Kemp, S.A., Dhar, M.S., Papa, G., Meng, B., Ferreira, I., Datir, R., Collier, D.A., Albecka, A., Singh, S., et al. (2021). SARS-CoV-2 B.1.617.2 Delta variant replication and immune evasion. Nature 599, 114–119. 10.1038/s41586-021-03944-y.

19. Liu, J., Liu, Y., Xia, H., Zou, J., Weaver, S.C., Swanson, K.A., Cai, H., Cutler, M., Cooper, D., Muik, A., et al. (2021). BNT162b2-elicited neutralization of B.1.617 and other SARS-CoV-2 variants. Nature 596, 273–275. 10.1038/s41586-021-03693-y.

20. Davis, C., Logan, N., Tyson, G., Orton, R., Harvey, W.T., Perkins, J.S., Mollett, G., Blacow, R.M., Consortium, C.-G.U., Peacock, T.P., et al. (2021). Reduced neutralisation of the Delta (B.1.617.2) SARS-CoV-2 variant of concern following vaccination. PLoS Pathog 17, e1010022. 10.1371/journal.ppat.1010022.

21. Cao, Y., Wang, J., Jian, F., Xiao, T., Song, W., Yisimayi, A., Huang, W., Li, Q., Wang, P., An, R., et al. (2022). Omicron escapes the majority of existing SARS-CoV-2 neutralizing antibodies. Nature 602, 657–663. 10.1038/s41586-021-04385-3.

22. Carreno, J.M., Alshammary, H., Tcheou, J., Singh, G., Raskin, A.J., Kawabata, H., Sominsky, L.A., Clark, J.J., Adelsberg, D.C., Bielak, D.A., et al. (2022). Activity of convalescent and vaccine serum against SARS-CoV-2 Omicron. Nature 602, 682–688. 10.1038/s41586-022-04399-5.

23. Menegale, F., Manica, M., Zardini, A., Guzzetta, G., Marziano, V., d’Andrea, V., Trentini, F., Ajelli, M., Poletti, P., and Merler, S. (2023). Evaluation of Waning of SARS-CoV-2 Vaccine-Induced Immunity: A Systematic Review and Meta-analysis. JAMA Netw Open 6, e2310650. 10.1001/jamanetworkopen.2023.10650.

24. Lu, L., Chen, L.L., Zhang, R.R., Tsang, O.T., Chan, J.M., Tam, A.R., Leung, W.S., Chik, T.S., Lau, D.P., Choi, C.Y., et al. (2022). Boosting of serum neutralizing activity against the Omicron variant among recovered COVID-19 patients by BNT162b2 and CoronaVac vaccines. EBioMedicine 79, 103986. 10.1016/j.ebiom.2022.103986.

25. Moreira, E.D., Jr., Kitchin, N., Xu, X., Dychter, S.S., Lockhart, S., Gurtman, A., Perez, J.L., Zerbini, C., Dever, M.E., Jennings, T.W., et al. (2022). Safety and Efficacy of a Third Dose of BNT162b2 Covid-19 Vaccine. N Engl J Med 386, 1910–1921. 10.1056/NEJMoa2200674.

26. Yan, K., Dumenil, T., Tang, B., Le, T.T., Bishop, C.R., Suhrbier, A., and Rawle, D.J. (2022). Evolution of ACE2-independent SARS-CoV-2 infection and mouse adaption after passage in cells expressing human and mouse ACE2. Virus Evol 8, veac063. 10.1093/ve/veac063.

27. Amarilla, A.A., Sng, J.D.J., Parry, R., Deerain, J.M., Potter, J.R., Setoh, Y.X., Rawle, D.J., Le, T.T., Modhiran, N., Wang, X., et al. (2021). A versatile reverse genetics platform for SARS-CoV-2 and other positive-strand RNA viruses. Nat Commun 12, 3431. 10.1038/s41467-021-23779-5.

28. Crawford, K.H.D., Eguia, R., Dingens, A.S., Loes, A.N., Malone, K.D., Wolf, C.R., Chu, H.Y., Tortorici, M.A., Veesler, D., Murphy, M., et al. (2020). Protocol and Reagents for Pseudotyping Lentiviral Particles with SARS-CoV-2 Spike Protein for Neutralization Assays. Viruses 12. 10.3390/v12050513.

29. Amarilla, A.A., Modhiran, N., Setoh, Y.X., Peng, N.Y.G., Sng, J.D.J., Liang, B., McMillan, C.L.D., Freney, M.E., Cheung, S.T.M., Chappell, K.J., et al. (2021). An Optimized High-Throughput Immuno-Plaque Assay for SARS-CoV-2. Front Microbiol 12, 625136. 10.3389/fmicb.2021.625136.

30. Hsieh, C.L., Goldsmith, J.A., Schaub, J.M., DiVenere, A.M., Kuo, H.C., Javanmardi, K., Le, K.C., Wrapp, D., Lee, A.G., Liu, Y., et al. (2020). Structure-based design of prefusion-stabilized SARS-CoV-2 spikes. Science 369, 1501–1505. 10.1126/science.abd0826.

31. Watterson, D., Wijesundara, D.K., Modhiran, N., Mordant, F.L., Li, Z., Avumegah, M.S., McMillan, C.L., Lackenby, J., Guilfoyle, K., van Amerongen, G., et al. (2021). Preclinical development of a molecular clamp-stabilised subunit vaccine for severe acute respiratory syndrome coronavirus 2. Clin Transl Immunology 10, e1269. 10.1002/cti2.1269.

32. McMillan, C.L.D., Choo, J.J.Y., Idris, A., Supramaniam, A., Modhiran, N., Amarilla, A.A., Isaacs, A., Cheung, S.T.M., Liang, B., Bielefeldt-Ohmann, H., et al. (2021). Complete protection by a single-dose skin patch-delivered SARS-CoV-2 spike vaccine. Sci Adv 7, eabj8065. 10.1126/sciadv.abj8065.

33. McMillan, C.L.D., Amarilla, A.A., Modhiran, N., Choo, J.J.Y., Azuar, A., Honeyman, K.E., Khromykh, A.A., Young, P.R., Watterson, D., and Muller, D.A. (2022). Skin-patch delivered subunit vaccine induces broadly neutralising antibodies against SARS-CoV-2 variants of concern. Vaccine 40, 4929–4932. 10.1016/j.vaccine.2022.07.013.

34. 34. ter Meulen, J., van den Brink, E.N., Poon, L.L., Marissen, W.E., Leung, C.S., Cox, F., Cheung, C.Y., Bakker, A.Q., Bogaards, J.A., van Deventer, E., et al. (2006). Human monoclonal antibody combination against SARS coronavirus: synergy and coverage of escape mutants. PLoS Med 3, e237. 10.1371/journal.pmed.0030237.

35. Isaacs, A., Amarilla, A.A., Aguado, J., Modhiran, N., Albornoz, E.A., Baradar, A.A., McMillan, C.L.D., Choo, J.J.Y., Idris, A., Supramaniam, A., et al. (2022). Nucleocapsid Specific Diagnostics for the Detection of Divergent SARS-CoV-2 Variants. Front Immunol 13, 926262. 10.3389/fimmu.2022.926262.

36. Liu, Y., Zhang, X., Liu, J., Xia, H., Zou, J., Muruato, A.E., Periasamy, S., Kurhade, C., Plante, J.A., Bopp, N.E., et al. (2022). A live-attenuated SARS-CoV-2 vaccine candidate with accessory protein deletions. Nat Commun 13, 4337. 10.1038/s41467-022-31930-z.

37. Chen, C., Nadeau, S., Yared, M., Voinov, P., Xie, N., Roemer, C., and Stadler, T. (2022). CoV-Spectrum: analysis of globally shared SARS-CoV-2 data to identify and characterize new variants. Bioinformatics 38, 1735–1737. 10.1093/bioinformatics/btab856.

38. Zou, J., Kurhade, C., Chang, H.C., Hu, Y., Meza, J.A., Beaver, D., Trinh, K., Omlid, J., Elghetany, B., Desai, R., et al. (2023). An Integrated Research-Clinical BSL-2 Platform for a Live SARS-CoV-2 Neutralization Assay. Viruses 15. 10.3390/v15091855.

39. Pajon, R., Doria-Rose, N.A., Shen, X., Schmidt, S.D., O’Dell, S., McDanal, C., Feng, W., Tong, J., Eaton, A., Maglinao, M., et al. (2022). SARS-CoV-2 Omicron Variant Neutralization after mRNA-1273 Booster Vaccination. N Engl J Med 386, 1088–1091. 10.1056/NEJMc2119912.

40. Wang, H., Xue, Q., Zhang, H., Yuan, G., Wang, X., Sheng, K., Li, C., Cai, J., Sun, Y., Zhao, J., et al. (2023). Neutralization against Omicron subvariants after BA.5/BF.7 breakthrough infection weakened as virus evolution and aging despite repeated prototype-based vaccination(1). Emerg Microbes Infect 12, 2249121. 10.1080/22221751.2023.2249121.

41. 41. Willett, B.J., Grove, J., MacLean, O.A., Wilkie, C., De Lorenzo, G., Furnon, W., Cantoni, D., Scott, S., Logan, N., Ashraf, S., et al. (2022). SARS-CoV-2 Omicron is an immune escape variant with an altered cell entry pathway. Nat Microbiol 7, 1161–1179. 10.1038/s41564-022-01143-7.

42. Mittal, A., Khattri, A., and Verma, V. (2022). Structural and antigenic variations in the spike protein of emerging SARS-CoV-2 variants. PLoS Pathog 18, e1010260. 10.1371/journal.ppat.1010260.

43. Wrapp, D., Wang, N., Corbett, K.S., Goldsmith, J.A., Hsieh, C.L., Abiona, O., Graham, B.S., and McLellan, J.S. (2020). Cryo-EM structure of the 2019-nCoV spike in the prefusion conformation. Science 367, 1260–1263. 10.1126/science.abb2507.

44. Lu, M., Chamblee, M., Zhang, Y., Ye, C., Dravid, P., Park, J.G., Mahesh, K.C., Trivedi, S., Murthy, S., Sharma, H., et al. (2022). SARS-CoV-2 prefusion spike protein stabilized by six rather than two prolines is more potent for inducing antibodies that neutralize viral variants of concern. Proc Natl Acad Sci U S A 119, e2110105119. 10.1073/pnas.2110105119.

45. Greaney, A.J., Loes, A.N., Crawford, K.H.D., Starr, T.N., Malone, K.D., Chu, H.Y., and Bloom, J.D. (2021). Comprehensive mapping of mutations in the SARS-CoV-2 receptor-binding domain that affect recognition by polyclonal human plasma antibodies. Cell Host Microbe 29, 463–476 e466. 10.1016/j.chom.2021.02.003.

46. Deng, X., Garcia-Knight, M.A., Khalid, M.M., Servellita, V., Wang, C., Morris, M.K., Sotomayor-Gonzalez, A., Glasner, D.R., Reyes, K.R., Gliwa, A.S., et al. (2021). Transmission, infectivity, and neutralization of a spike L452R SARS-CoV-2 variant. Cell 184, 3426–3437 e3428. 10.1016/j.cell.2021.04.025.

47. Neerukonda, S.N., Vassell, R., Lusvarghi, S., Wang, R., Echegaray, F., Bentley, L., Eakin, A.E., Erlandson, K.J., Katzelnick, L.C., Weiss, C.D., and Wang, W. (2021). SARS-CoV-2 Delta Variant Displays Moderate Resistance to Neutralizing Antibodies and Spike Protein Properties of Higher Soluble ACE2 Sensitivity, Enhanced Cleavage and Fusogenic Activity. Viruses 13. 10.3390/v13122485.

48. Xie, X., Liu, Y., Liu, J., Zhang, X., Zou, J., Fontes-Garfias, C.R., Xia, H., Swanson, K.A., Cutler, M., Cooper, D., et al. (2021). Neutralization of SARS-CoV-2 spike 69/70 deletion, E484K and N501Y variants by BNT162b2 vaccine-elicited sera. Nat Med 27, 620–621. 10.1038/s41591-021-01270-4.

49. Wang, Z., Schmidt, F., Weisblum, Y., Muecksch, F., Barnes, C.O., Finkin, S., Schaefer-Babajew, D., Cipolla, M., Gaebler, C., Lieberman, J.A., et al. (2021). mRNA vaccine-elicited antibodies to SARS-CoV-2 and circulating variants. Nature 592, 616–622. 10.1038/s41586-021-03324-6.

50. Qu, P., Evans, J.P., Zheng, Y.M., Carlin, C., Saif, L.J., Oltz, E.M., Xu, K., Gumina, R.J., and Liu, S.L. (2022). Evasion of neutralizing antibody responses by the SARS-CoV-2 BA.2.75 variant. Cell Host Microbe 30, 1518–1526 e1514. 10.1016/j.chom.2022.09.015.

51. Cox, M., Peacock, T.P., Harvey, W.T., Hughes, J., Wright, D.W., Consortium, C.-G.U., Willett, B.J., Thomson, E., Gupta, R.K., Peacock, S.J., et al. (2023). SARS-CoV-2 variant evasion of monoclonal antibodies based on in vitro studies. Nat Rev Microbiol 21, 112–124. 10.1038/s41579-022-00809-7.

52. Chen, D.Y., Chin, C.V., Kenney, D., Tavares, A.H., Khan, N., Conway, H.L., Liu, G., Choudhary, M.C., Gertje, H.P., O’Connell, A.K., et al. (2023). Spike and nsp6 are key determinants of SARS-CoV-2 Omicron BA.1 attenuation. Nature 615, 143–150. 10.1038/s41586-023-05697-2.

53. Demonbreun, A.R., Sancilio, A., Velez, M.P., Ryan, D.T., Saber, R., Vaught, L.A., Reiser, N.L., Hsieh, R.R., D’Aquila, R.T., Mustanski, B., et al. (2021). Comparison of IgG and neutralizing antibody responses after one or two doses of COVID-19 mRNA vaccine in previously infected and uninfected individuals. EClinicalMedicine 38, 101018. 10.1016/j.eclinm.2021.101018.

54. Chen, W., Zhang, L., Li, J., Bai, S., Wang, Y., Zhang, B., Zheng, Q., Chen, M., Zhao, W., and Wu, J. (2022). The kinetics of IgG subclasses and contributions to neutralizing activity against SARS-CoV-2 wild-type strain and variants in healthy adults immunized with inactivated vaccine. Immunology 167, 221–232. 10.1111/imm.13531.

55. Winichakoon, P., Wipasa, J., Chawansuntati, K., Salee, P., Sudjaritruk, T., Yasri, S., Khamwan, C., Peerakam, R., Dankai, D., and Chaiwarith, R. (2023). Diagnostic performance between in-house and commercial SARS-CoV-2 serological immunoassays including binding-specific antibody and surrogate virus neutralization test (sVNT). Sci Rep 13, 34. 10.1038/s41598-022-26202-1.

56. Takheaw, N., Liwsrisakun, C., Chaiwong, W., Laopajon, W., Pata, S., Inchai, J., Duangjit, P., Pothirat, C., Bumroongkit, C., Deesomchok, A., et al. (2022). Correlation Analysis of Anti-SARS-CoV-2 RBD IgG and Neutralizing Antibody against SARS-CoV-2 Omicron Variants after Vaccination. Diagnostics (Basel) 12. 10.3390/diagnostics12061315.

57. Bewley, K.R., Coombes, N.S., Gagnon, L., McInroy, L., Baker, N., Shaik, I., St-Jean, J.R., St-Amant, N., Buttigieg, K.R., Humphries, H.E., et al. (2021). Quantification of SARS-CoV-2 neutralizing antibody by wild-type plaque reduction neutralization, microneutralization and pseudotyped virus neutralization assays. Nat Protoc 16, 3114–3140. 10.1038/s41596-021-00536-y.

